# Cell Type-Specific Chromatin Accessibility Analysis in the Mouse and Human Brain

**DOI:** 10.1101/2020.07.04.188094

**Authors:** Devin Rocks, Ivana Jaric, Lydia Tesfa, John M. Greally, Masako Suzuki, Marija Kundakovic

## Abstract

The Assay for Transposase Accessible Chromatin by sequencing (ATAC-seq) is becoming increasingly popular in the neuroscience field where chromatin regulation is thought to be involved in neurodevelopment, activity-dependent gene regulation, hormonal and environmental responses, and the pathophysiology of neuropsychiatric disorders. The advantages of using this assay include a small amount of material needed, relatively simple and fast protocol, and the ability to capture a range of gene regulatory elements with a single assay. However, with increasing interest in chromatin research, it is an imperative to have feasible, reliable assays that are compatible with a range of neuroscience study designs in both animals and humans. Here we tested three different protocols for neuronal chromatin accessibility analysis, including a varying brain tissue freezing method followed by fluorescent-activated nuclei sorting (FANS) and the ATAC-seq analysis. Our study shows that the cryopreservation method impacts the number of open chromatin regions that can be identified from frozen brain tissue using the cell-type specific ATAC-seq assay. However, we show that all three protocols generate consistent and robust data and enable the identification of functional regulatory elements, promoters and enhancers, in neuronal cells. Our study also implies that the broad biological interpretation of chromatin accessibility data is not significantly affected by the freezing condition. In comparison to the mouse brain analysis, we reveal the additional challenges of doing chromatin analysis on *post mortem* human brain tissue. However, we also show that these studies are revealing important cell type-specific information about gene regulation in the human brain. Overall, the ATAC-seq coupled with FANS is a powerful method to capture cell-type specific chromatin accessibility information in the mouse and human brain. Our study provides alternative brain preservation methods that generate high quality ATAC-seq data while fitting in different study designs, and further encourages the use of this method to uncover the role of epigenetic (dys)regulation in healthy and malfunctioning brain.

## INTRODUCTION

Chromatin, the dynamic nuclear complex of genomic DNA and its associated proteins (mostly histones), controls DNA packaging and accessibility of DNA regulatory elements in eukaryotic cells^1^. Chromatin modifications, including DNA and histone modifications, confer local chromatin states that can be either permissive (“open” chromatin) or non-permissive (“closed” chromatin) to the binding of transcriptional regulators. In addition, a special class of transcription factors (TFs), pioneer TFs, are able to bind “closed” chromatin, trigger its opening, and directly make it competent for other factors to bind^2^. Control of chromatin organization is, therefore, a major regulatory mechanism controlling gene expression^3^ and has been extensively studied across biomedical fields.

In neuroscience, chromatin regulation has been shown to be critical for transcriptional programs underlying brain development, neuronal maturation, and plasticity in mature mammalian brain^4,5^. In neurons, chromatin mediates responses to both endogenous (e.g. hormones, neurotransmitters)^6^ and exogenous (e.g. drugs, stress)^7^ factors contributing to neuronal-specific information processing and neuroplasticity. In addition, chromatin alterations may mediate long-term effects of early life stressors on neuronal and glial gene expression, brain structure, and behavior^8^. Finally, there is a great interest in revealing how gene regulatory landscapes may be altered in major neuropsychiatric disorders to advance our understanding of the molecular underpinnings of mental illness^9^.

With all this interest in chromatin research, it is an imperative to have feasible, reliable assays that are compatible with a range of neuroscience study designs in both animals and humans. The chromatin immunoprecipitation using sequencing (ChIP-seq) analysis of histone modifications used to be a gold standard for chromatin research^10^, with specific histone marks used as proxies for the chromatin state around regulatory regions^11-14^. However, histone ChIP-seq posed many limitations for neuroscientists including: the need for a high amount of starting material; complicated, multiday protocols; and assays on single histone marks providing limited information about gene regulatory elements across the genome^15^.

More recently, chromatin analysis using the assay for transposase-accessible chromatin using sequencing (ATAC-seq) was introduced^16^, and has become increasingly popular in the neuroscience field. The ATAC-seq method requires only 50,000 cells or less; involves an easy and quick protocol^17^; and the resulting data capture regulatory elements more comprehensively^1^, replacing the testing for multiple histone modifications or other chromatin components. The original method seemed limited to the analyses of fresh cells and tissues, due to the disruptive effect of freezing on chromatin structure and integrity^16,18,19^. However, optimized assays such as omni-ATAC-seq^20^ have been developed and several studies using ATAC-seq have now been performed to interrogate chromatin organization from frozen mouse and human brain tissue^6,21-26^.

While working with frozen brain tissue is almost inevitable in neuroscience studies, the cryopreservation of the tissue can be done using different methods, and the data comparing different experimental conditions is missing. In this study, we examined the efficacy and reliability of a cell-type specific ATAC-seq assay^6^ across three different experimental conditions. We selected scenarios that are compatible with the study designs that we and others typically use in our research, including flash-freezing of brain tissue using either liquid nitrogen^6^ or dry ice cooled hexane^27,28^, and compared them with a slow freezing protocol using a cryoprotective medium^18^. For an additional comparison, we included the analysis of human *post mortem* brain tissue using the same ATAC-seq protocol. Together, our data show that the analyzed methods are highly consistent, particularly in detecting robust ATAC-seq signals; however, more controlled freezing conditions allow detection of a higher number of open chromatin regions. Our results encourage further use of the ATAC-seq method for cell-type specific chromatin profiling of the mouse and human brain tissue and provide guidelines for alternative study designs and quality control check points.

## RESULTS

### Mouse Study Design

To assess the impact of the brain tissue cryopreservation method on chromatin accessibility profiles in neuronal cells, we included three freezing conditions in our mouse study: 1) controlled slow freezing in a cryoprotective medium (MED); 2) rapid freezing in the dry ice cooled hexane (HEX); and 3) immediate flash freezing using liquid nitrogen (LN2) (**Figure 1A**). For this study, we assessed the frontal cortex of adult male mice, using two biological replicates for each condition. We included male mice only to avoid the confounding biological effect of sex and estrous cycle on chromatin organization^6^. From each condition, nuclei were prepared using the sucrose gradient method, and the neuronal (NeuN+) nuclei fraction was purified by fluorescence-activated nuclei sorting (FANS), using an antibody against the neuronal nuclear marker NeuN^15^. Chromatin accessibility analysis of NeuN+ nuclei was performed using ATAC-seq^16^ (**Figure 1A**). In this assay, a hyperactive Tn5 transposase enzyme cuts DNA and inserts sequencing adapters into open chromatin regions. These open or accessible DNA regions are then enriched using PCR amplification and identified, genome-wide, using the next generation sequencing and bioinformatic analysis. Each open chromatin region (ATAC-seq peak) is identified using a peak caller, and reproducible peaks are recovered by the Irreproducible Discovery Rate (IDR) analysis, as per the Encyclopedia of DNA Element (ENCODE) guidelines^10^. We performed a comprehensive quality control and analysis of the ATAC-seq data, with more than 40 million mapped paired-end reads produced from each condition (**Suppl. Table 1**).

**Figure 1.**
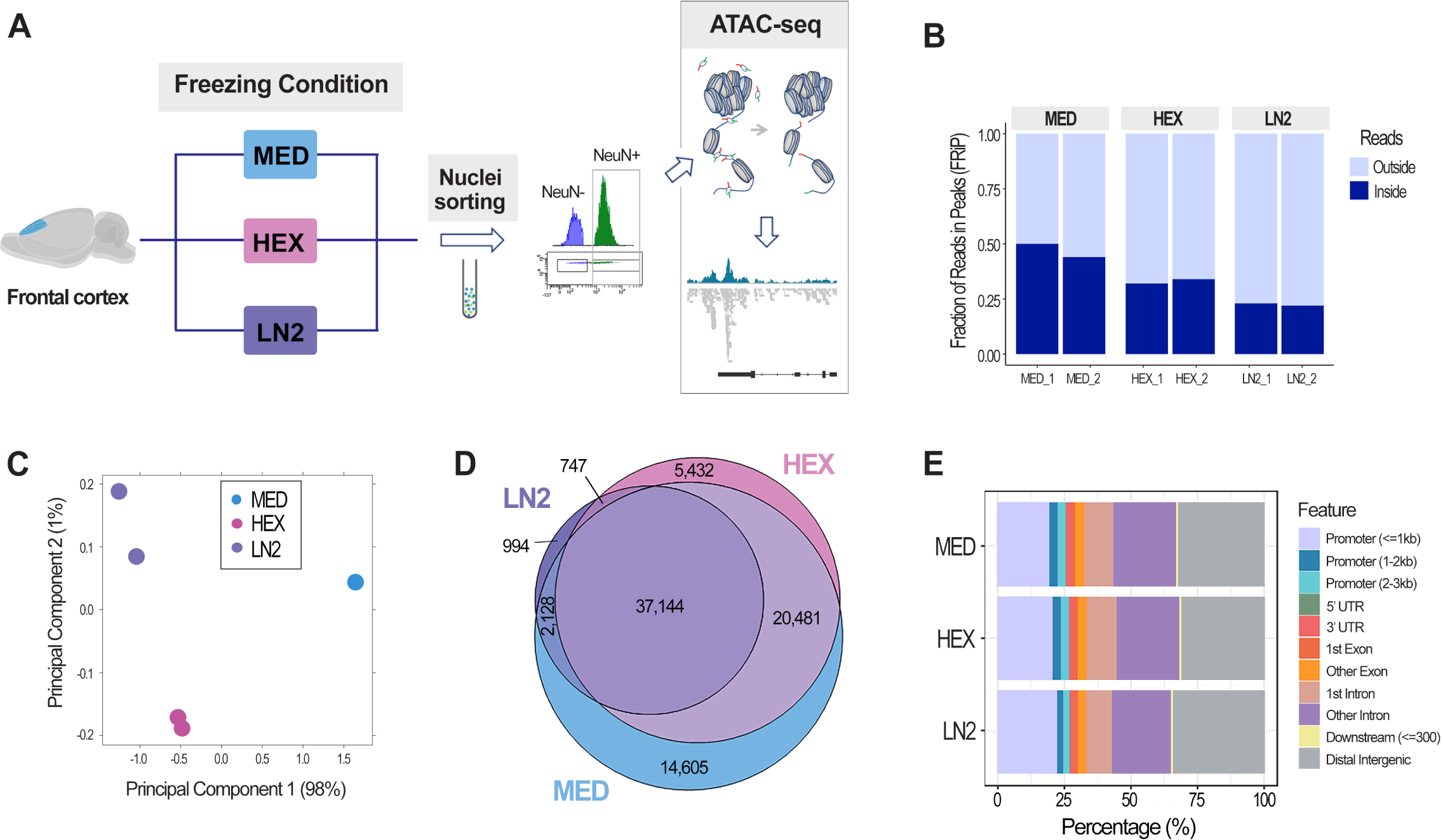
Chromatin accessibility analysis using cell-type specific ATAC-seq assay in the mouse brain. A. Scheme of the ATAC-seq assay; three freezing conditions were used on dissected mouse frontal cortex: slow freezing in a cryoprotective medium (MED), and flash freezing in either dry ice cooled hexane (HEX) or liquid nitrogen (LN2). Neuronal (NeuN+) nuclei were purified with FANS and used for the analysis. B. FRiP score shows fraction of reads in peaks as a QC measure of enrichment in all three conditions: MED, HEX, and LN2. C. Principal component analysis of the ATAC-seq duplicates for each condition. D. Venn diagrams show the number of open chromatin regions (ATAC-seq peaks) in each condition following the IDR analysis and their overlap. E. Distribution of the reproducible (IDR) peaks across the genome in each condition.

### The effect of the cryopreservation method on ATAC-seq signal enrichment, peak number, and distribution

The FRiP (Fraction of Reads in Peaks) score is an important quality control metric representing the fraction of all mapped reads that fall into the called ATAC-seq peaks, as opposed to random reads or non-specific background^10^ (**Figure 1B**). Typically, FRiP values correlate positively with the number of called regions and values greater than 0.2 are considered acceptable by the ENCODE standards (https://www.encodeproject.org/atac-seq/). While all three conditions passed the ENCODE standards, the FRiP scores were highest for the duplicates in the MED condition (0.44 and 0.50), followed by those in the HEX condition (0.32 and 0.34), and finally by the duplicates in the LN2 condition (0.22 and 0.23) (**Figure 1B**). Based on a principal component analysis, the biological replicates belonging to the three examined conditions also clearly clustered by freezing condition (**Figure 1C**). After performing the IDR analysis, which retains only high confidence peaks in each condition (**Suppl. Table 2**), we performed differential chromatin accessibility analysis between the groups (**Fig. 1D**). This analysis showed that, among all open chromatin regions or peaks (81,531), the highest number of peaks was found in the MED condition (74,358 or 91.20%), then in the HEX condition (63,804 or 78.26%), and the lowest number in the LN2 condition (41,013 or 50.30%) (**Fig. 1D**). While we note a significant difference in the number of peaks between the three conditions, there was also a striking overlap between the three groups. In fact, the LN2 group, the group with the lowest number of peaks, shared 90.57% (37,144 out of 41,013) of accessible chromatin regions with the other two groups (**Fig. 1D**). In addition, peak distribution across the genome looked comparable between the three groups, with around 25% of peaks located within the putative promoter regions and more than 30% of peaks located in the distal intergenic regions (**Fig. 1E**).

Together, these results show that, although there are notable differences in FRiP scores and number of peaks, all three conditions passed rigorous ATAC-seq quality standards, showing a significant overlap in the accessible chromatin regions detected.

### ATAC-seq peak height distribution

To further examine differences in the ATAC-seq data, we looked at the peak height distribution for each condition (**Fig. 2A**). Consistent with the FRiP scores (**Fig. 1B**), we found that the peak height was generally highest in the MED condition, followed by the HEX group and, finally, the LN2 condition (**Fig. 2A**). We then looked into the MED-specific peaks, the peaks that were called only in the MED group but not in the HEX or LN2 group (14,605 or 17.91% of all peaks, **Fig. 1D**). As expected, the reads corresponding to the MED-specific peaks were enriched in the MED condition (**Fig. 2B**); however, while not passing the threshold for peak calling, the corresponding reads were also detected in the HEX and LN2 groups (**Fig. 2B**). Finally, we compared the peak height of the MED-specific peaks with the average peak height in the MED condition (**Fig. 2C**). These data showed that the MED-specific peaks were the smallest peaks in the MED condition (**Fig. 2C**), which suggested that slow freezing improves preservation of weak peaks or uncovers the regions with low enrichment of open chromatin, but have no significant effect on strong ATAC-seq peaks.

**Figure 2.**
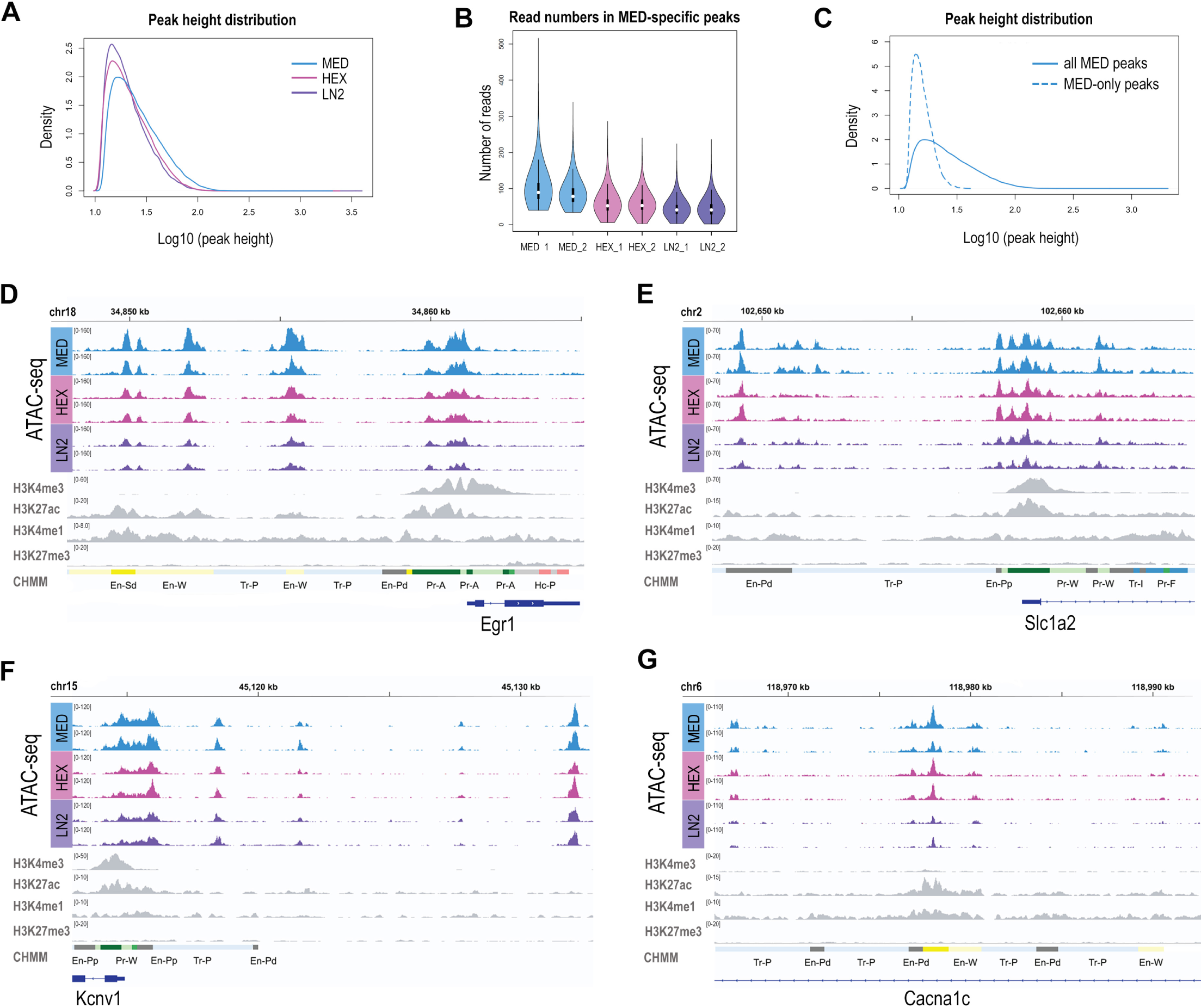
Chromatin signatures in neuronal cells of the mouse brain. ATAC-seq peak height (A) and Read number in MED-specific peaks (B) for MED, HEX, and LN2 condition. C. Average peak height in all MED peaks vs. MED-only peaks. Chromatin accessibility (ATAC-seq) profiles of *Egr1* (D), *Slc1a2* (E), *Kcnv1* (F), and *Cacna1c* (G) in mouse cortical neuronal cells, shown in duplicates for each condition. Histone modification ChIP-seq tracks (H3K4me3, H3K27ac, H3K4me1, H3K27me3) for all four genes are derived from postnatal 0 day mouse forebrain bulk tissue, generated by the Mouse ENCODE project. MED neuronal nuclei, blue; HEX neuronal nuclei, pink; LN2 neuronal nuclei, purple; ENCODE bulk brain tissue, gray.

### ATAC-seq signal in neuronal-relevant genes

To further compare the ATAC-seq signal among three experimental conditions, we explored the ATAC-seq peaks surrounding four specific genes of relevance to neuronal function (**Fig. 2D-G**). The early growth response (*Egr1*) gene was of interest as a neuronal activity-dependent, immediate early gene encoding transcription factor^29^. *Slc1a2* encodes the glutamate transporter 1, critical for the removal of excess glutamate from the synapse^30^. *Kcnv1* is a gene encoding a potassium voltage-gated channel relevant to neuronal excitability^31^. Finally, *Cacna1c* is a candidate psychiatric risk gene encoding a voltage-gated L-type calcium channel^32^. We looked at our ATAC-seq data in parallel with publicly available Mouse ENCODE ChIP-seq data^33,34^ on four histone modifications that are, in combination, used as proxies for: active promoters (H3K4me3/H3K27ac), active enhancers (H3K4me1/H3K27ac), and poised enhancers (H3K4me1/H3K27me3)^11-13^. We also included the ENCODE chromatin state (CHMM) annotations^33^ generated by the chromHMM method^35^, which integrates ChIP-seq data from 8 histone modifications in order to predict functional regulatory elements, promoters and enhancers. All used ENCODE data were generated from the postnatal day 0 (P0) mouse bulk forebrain tissue.

Although the height of the ATAC-seq peaks varied (MED>HEX>LN2), in all three conditions, we found a comparable peak pattern for all four genes (**Fig. 2D-G)**. The *Egr1* gene showed a clear ATAC-seq signal adjacent to the transcription start site (TSS), typical of an active promoter (**Fig. 2D**). These data were consistent with strong H3K4me3 and H3K27ac signals and the CHMM prediction of an active promoter in the ENCODE data (**Fig. 2D**). For Egr1, we also detected three putative enhancers within 10 kb upstream of the TSS, which were associated with ATAC-seq peaks in all three conditions and corresponded with the combined H3K27ac/H3K4me1 ENCODE signal and the enhancer prediction by the chromHMM algorithm (**Fig. 2D**).

For *Slc1a2*, the broad ATAC-seq peak around the TSS corresponded to both predicted promoter (combined H3K4me3/H3K27ac signal) and small proximal enhancer (H3K27ac/H3K4me1) from the ENCODE data (**Fig. 2E**). We also detected an ATAC-seq peak within 10 kb upstream of the TSS, consistent with a distal enhancer (corresponded to the H3K27ac/H3K4me1 signal) (**Fig. 2E**).

For *Kcnv1*, similar to *Slc1a2*, the broad ATAC-seq peak around the TSS corresponded to both predicted promoter (combined H3K4me3/H3K27ac signal) and proximal enhancer (H3K27ac/H3K4me1) (**Fig. 2F**). However, our ATAC-seq data from all three conditions uncovered at least three putative enhancers within 15 kb upstream of the TSS which were not predicted by the ENCODE histone modification data (**Fig. 2F**). Since ENCODE data were generated from P0 forebrain tissue, these regulatory elements are likely to be active in a spatially- and temporally-specific manner in the mouse brain.

Finally, we identified an active intronic enhancer in the *Cacna1c* gene; the ATAC-seq peak found in all three conditions corresponded with the combined H3K27ac/H3K4me1 signal and the predicted CHMM enhancer in the ENCODE data (**Fig. 2G**).

Together, these data show that, although the strength of the ATAC-seq signal varies with the freezing condition, all three conditions enable the identification of functional regulatory regions, both promoters and enhancers.

### Gene Ontology and Pathway analysis of the ATAC-seq data

Considering that the number of peaks was significantly different between the three conditions, we wanted to further examine whether the different tissue preservation methods may affect the more general biological interpretation of the ATAC-seq data. Therefore, we performed the Gene Ontology (GO) and KEGG pathway analyses to gain an insight into biological functions of genes proximal to the regions found to have open chromatin in the three conditions. For the GO biological process, the top enriched terms showed a very similar pattern across the three conditions (**Fig. 3A, Suppl. Table 3**), and were broadly important for cellular function including nucleic acid and protein-related processes, mitochondrial and chromatin organization, and signaling transduction. The GO cellular component analysis revealed synaptic, neuronal-specific terms as well as more general cellular components with, again, a striking similarity between the three conditions (**Fig. 3A, Suppl. Table 3**). Finally, the KEGG pathway analysis showed comparable enrichment of genes relevant to protein processing, axon guidance, MAPK and hormone-related signaling pathways, among others, in all three conditions (**Fig. 3A, Suppl. Table 4**). Overall, the results of gene enrichment analyses were comparable across the three conditions, indicating that the broad biological interpretation of the data is not significantly affected by the freezing condition.

**Figure 3.**
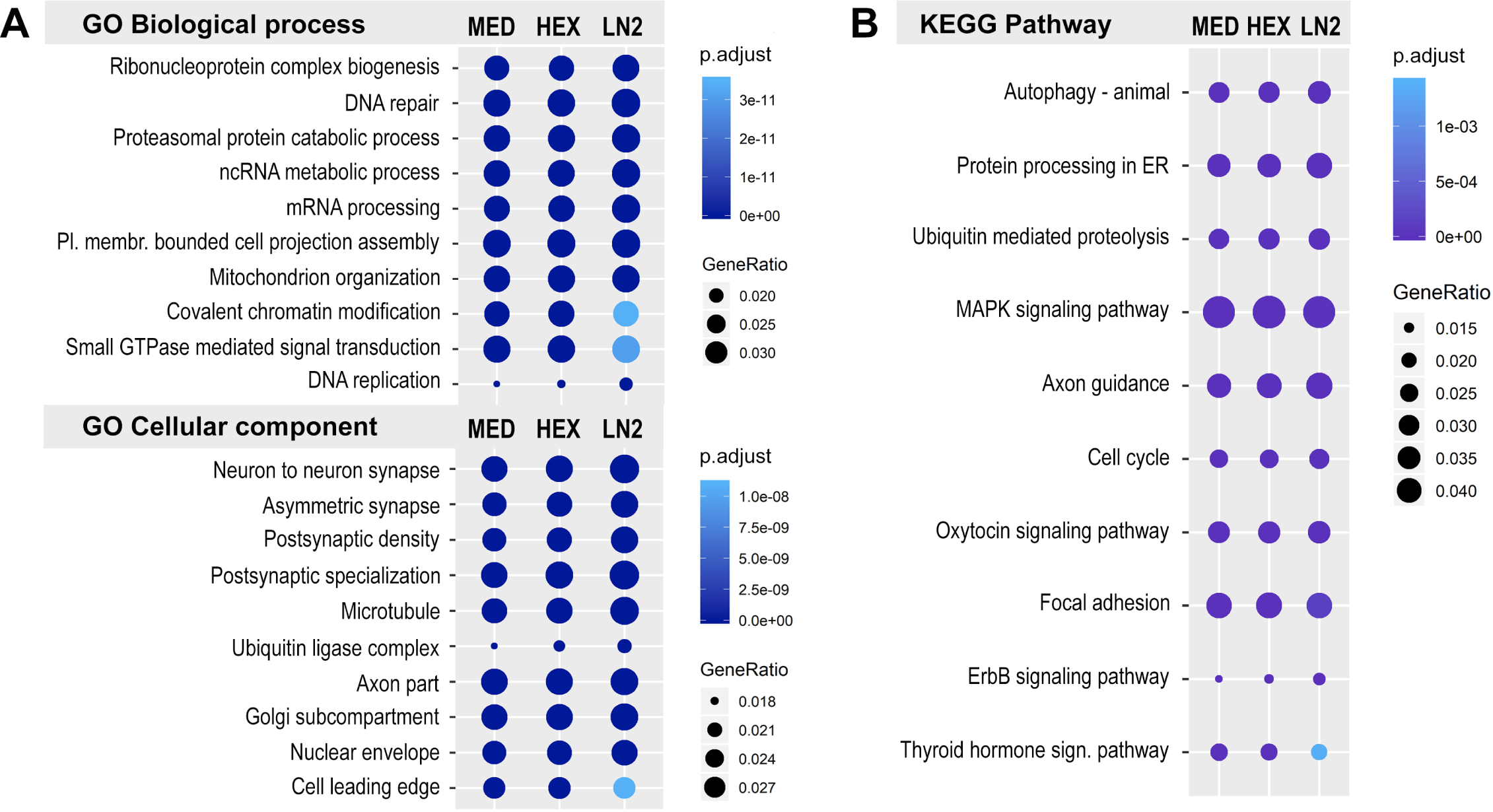
Gene ontology and KEGG pathway analysis of the mouse ATAC-seq data. A. Top ten GO terms for Biological process and Cellular component, and B. Top ten KEGG pathways in MED, HEX, and LN2 condition, with the MED condition used as a reference. Colors indicate adjusted p values and dot size corresponds to gene ratio.

### Motif analysis

Considering that the highest number of peaks was called in the MED condition, followed by the HEX condition and then the LN2 condition, we also wanted to examine whether there was any sequence bias in the ATAC-seq data that may be dependent on the freezing method. Therefore, we performed the HOMER motif analysis to assess the enrichment of transcription factor (TF) binding sites in the open chromatin regions recovered after each condition (**Fig. 4, Suppl. Table 5**). In all three conditions, we found similar enrichment of TF motifs including: the top enriched motif for the major higher-order chromatin organization protein CTCF^36^; motifs for the activator protein 1 (AP1) family of TFs involved in neuronal activity-dependent gene regulation^37^; motifs for the myocyte enhancer factor 2 (MEF2) family of TFs involved in synaptic plasticity^38,39^, and neural basic-helix-loop-helix (bHLH) family of TFs^40^ (**Fig. 4**). Therefore, our motif analysis indicates that no significant sequence bias has been observed in the ATAC-seq data generated using different freezing conditions.

**Figure 4.**
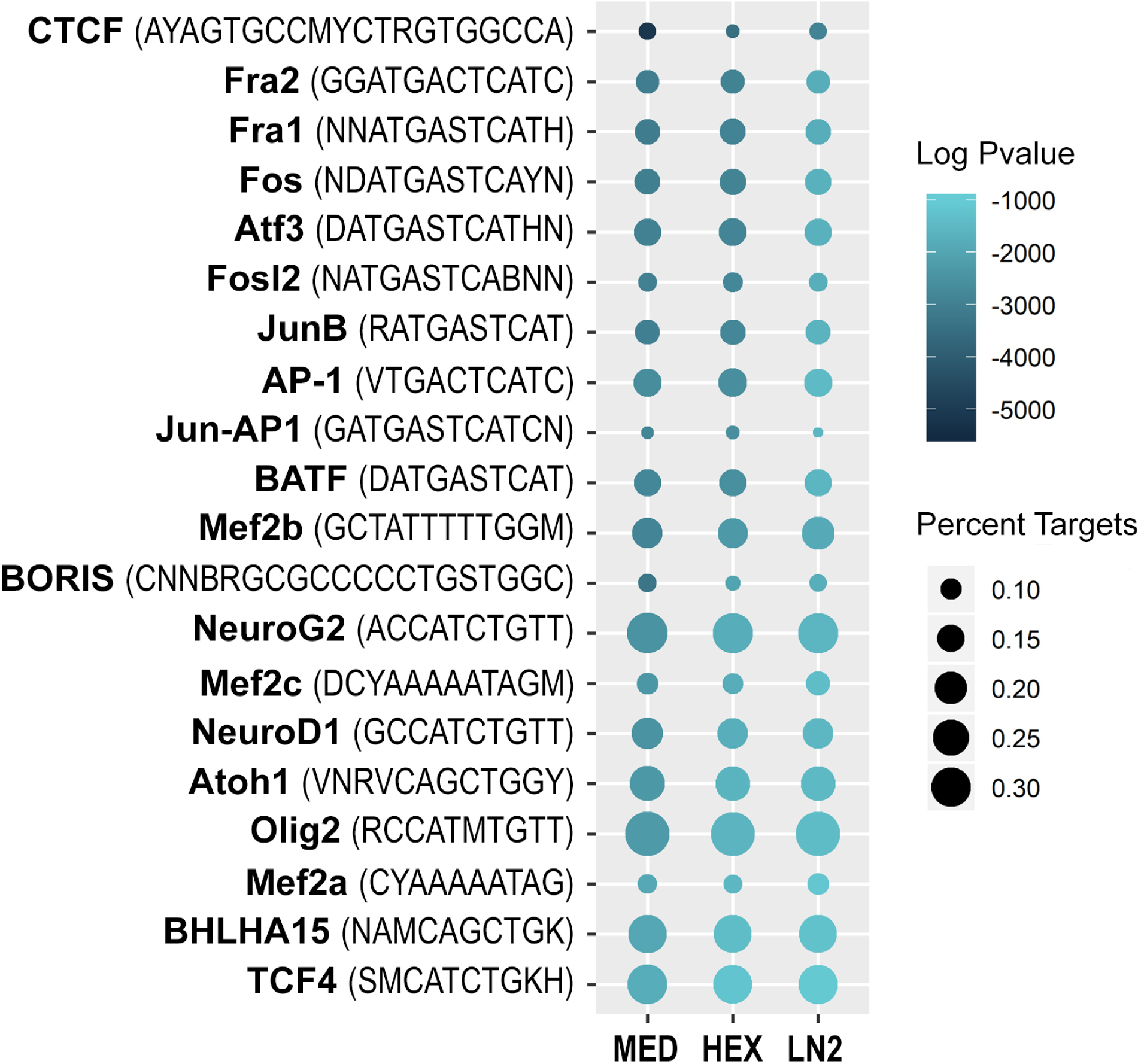
Motif analysis of the mouse ATAC-seq data. Dotplot shows top 20 transcription factor binding motifs in MED, HEX, and LN2 condition, with the MED condition used as a reference. Colors indicate log adjusted P values and dot size corresponds to percent target.

### ATAC-seq signal in neuronal nuclei of postmortem human brain

We also performed the ATAC-seq analysis on neuronal (NeuN+) nuclei isolated from two *post mortem* human (male) brains, specifically from the adult anterior cingulate cortex (ACC) (**Fig. 5A, Suppl. Table 6**), with the same procedure described above. While in terms of tissue preservation method the *post mortem* human brain tissue is closest to the LN2 mouse condition, the quality of human brain tissue is largely dependent on the *post mortem* interval (PMI, or the amount of time elapsed between a subject’s death and processing of tissue) which may have an impact on chromatin integrity and organization^41^. It was, therefore, not surprising that FRiP scores for the ATAC-seq data from human neuronal nuclei were lower (0.11 and 0.10, **Fig. 5A**) than those from the mouse neuronal nuclei (**Fig. 1B**), and were below the standard set by the ENCODE project. When we called the ATAC-seq peaks from neuronal nuclei of each human brain (A and B), we found that the peak distribution was similar to that seen in mouse neurons, with around 25% of the peaks located in the promoter region (**Fig. 5B**). However, a lower percentage of peaks were located in the distal intergenic regions, and a higher percentage of peaks were located in intronic regions, possibly indicating more intronically-located enhancers in human neurons compared to those of mice (**Fig. 5B**). Importantly, when we used the IDR method to select high confidence peaks, we found that the peak distribution changed significantly (A/B IDR, **Fig. 5B**). The majority of the peaks that survived the IDR method were located in the promoters (>50% of all peaks), while the percentage of intronic and distal intergenic peaks, likely corresponding to enhancers, fell significantly (**Fig. 5B**).

**Figure 5.**
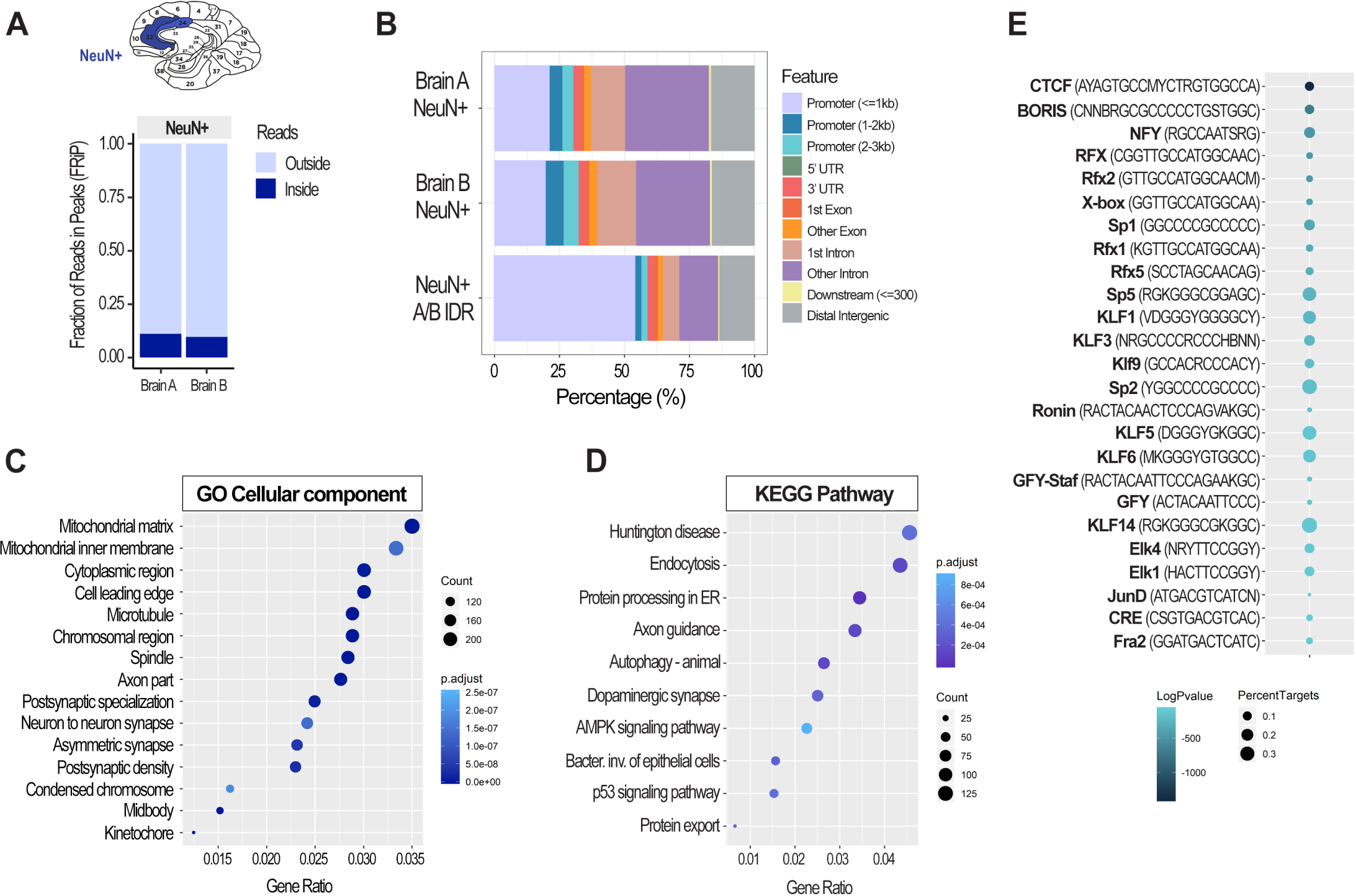
Chromatin accessibility analysis using cell-type specific ATAC-seq assay in postmortem human brain. FRiP scores (A) and genomic peak distribution (B) of the ATAC-seq data generated from neuronal (NeuN+) nuclei of the two postmortem human anterior cingulate cortices (Brodmann areas 24/32; Brains A and B). A/B IDR peaks, high confidence peaks generated by the IDR analysis, were then used for the GO Cellular component (C) and KEGG pathway (D) analyses. Colors indicate adjusted p values and dot size corresponds to gene count. Top 15 GO and top 10 KEGG pathways are shown. Motif analysis of the ATAC-seq data (E) shows 25 top transcription factor binding motifs. Colors indicate log adjusted P values and dot size corresponds to percent target.

We then took the high confidence peaks and performed the enrichment analyses including the GO, KEGG pathway, and motif analysis (**Fig. 5C-E**). The top GO cellular component terms for human cortical neurons (**Fig. 5C**) were comparable to those found in the mouse study (**Fig. 3A**) and included synaptic and neuronal-specific terms as well as more general cellular components (**Fig. 5C, Suppl. Table 7**). The KEGG pathway analysis revealed the top pathways that were shared with the mouse study including autophagy, protein processing in ER, axon guidance, and AMPK signaling pathway (**Fig. 5D**). In human neurons, additional neuronal-specific pathways involved Huntington disease-related genes and the dopaminergic synapse involving genes important for cognition, motivation, award, and motor control (**Fig. 5D, Suppl. Table 8**). In terms of the motif analysis, the top enriched TF motif was that for the chromatin organizational protein CTCF (**Fig. 5E**), which corresponded to the mouse data (**Fig. 4**). Among the top motifs, there were also motifs that were specific for human neurons including motifs for the regulatory factor X (RFX) and the Krüppel-like factor and specificity protein (KLF/SP) families of TFs (**Fig. 5E, Suppl. Table 9**). We also found that motifs for immediate early gene-encoded TFs were among top enriched motifs in humans (**Fig. 5E**), similar to that found in the mouse motif analysis (**Fig. 4**).

In summary, although the quality control analysis indicates that the ATAC-seq data generated from human *post mortem* brain tissue is of a lower quality than the data from the properly frozen mouse brain tissue, the bioinformatics analysis of the human data show expected enrichment of genes, pathways, and TF motifs relevant to neuronal function.

### The epigenomic signatures of neuronal and non-neuronal cells in postmortem human brain

Finally, we decided to focus on the human chromatin accessibility data related to two specific genes, *EGR1* and glial fibrillary acidic protein (*GFAP*). *EGR1* is expressed in the human brain with the assumed function of an activity-dependent immediate early gene^42^, similar to that found in the mouse brain. Therefore, we wanted to make a comparison of the human neuronal ATAC-seq data (**Fig. 6A**) with the corresponding mouse neuronal data (**Fig. 2D**). On the other hand, GFAP is a widely used glial protein marker^43^ specifically expressed by astrocytes and not found in neurons. Cell type-specific *GFAP* expression is established during brain development and involves DNA demethylation of the regulatory region in astrocytes while all non-expressing cells, including neurons, show high levels of DNA methylation within the same region^44,45^. In order to further assess the quality of the human chromatin accessibility data, we compared the ATAC-seq signal in the vicinity of the *GFAP* gene in neuronal (NeuN+) and non-neuronal (NeuN-) cells from the same postmortem human brain, and integrated these data with the corresponding DNA methylation data (**Fig. 6B**). For both genes, we explored our data in parallel with four histone modification ChIP-seq datasets (H3K4me3, H3K27ac, H3K4me1, H3K27me3) derived from the adult human male cingulate gyrus bulk tissue as part of the NIH Roadmap Epigenomics project^46^.

**Figure 6.**
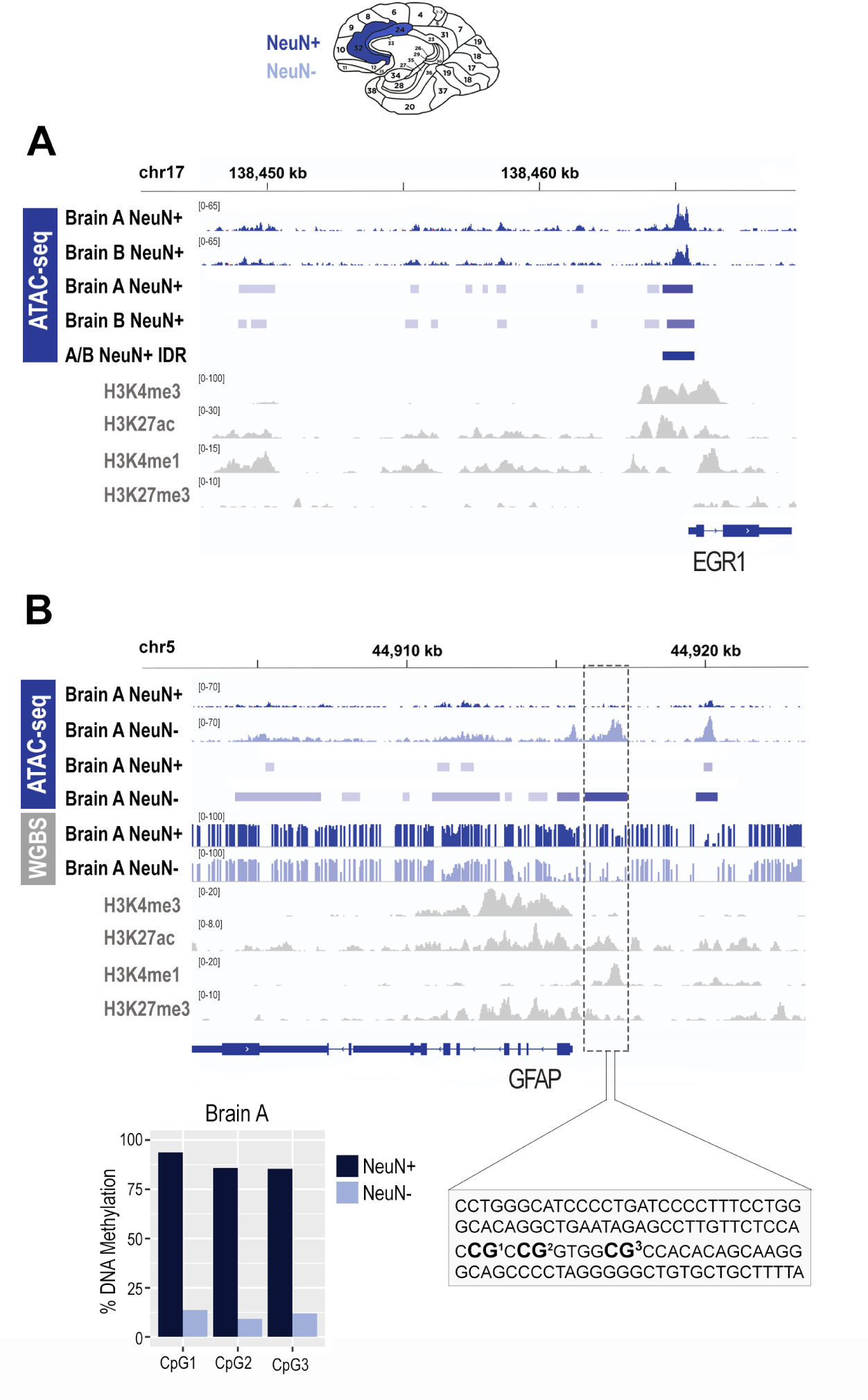
Epigenomic signatures of neuronal and non-neuronal cells in postmortem human brain. A. Chromatin accessibility (ATAC-seq) profile of the *EGR1* gene in neuronal (NeuN+) cells of the postmortem human anterior cingulate cortex; bam and bed files are shown for each brain separately (A and B); the A/B IDR track represents a bed file of high confidence peaks generated by the IDR analysis. B. Chromatin accessibility (ATAC-seq) and DNA methylation (WGBS) profile of the *GFAP* gene in neuronal (NeuN+) and non-neuronal (NeuN-) cells of the postmortem human anterior cingulate cortex; bam, bed, and bedgraph files are shown for both cell types; shown is also a focused DNA methylation analysis of the three CpG sites (bar graph) located in the proximal enhancer (dashed rectangle) using bisulfite pyrosequencing. Histone modification ChIP-seq tracks (H3K4me3, H3K27ac, H3K4me1, H3K27me3) for both genes are derived from the cingulate gyrus bulk tissue, generated by the NIH Roadmap Epigenomics project. Neuronal (NeuN+) nuclei, dark blue; Non-neuronal (NeuN-) nuclei, light blue; NIH Roadmap Epigenomics bulk brain tissue, gray.

For *EGR1* in neuronal (NeuN+) nuclei, we observed a strong ATAC-seq signal (open chromatin) close to the TSS, which is consistent with an active promoter, and consistent with the histone modification data showing strong enrichment of H3K4me3 and H3K27ac within the same region (**Fig. 6A**). When we looked at the region within 15 kb upstream of the TSS, consistent with the histone modification data (H3K27ac/H3K4me1), we see several ATAC-seq peaks being called in neuronal nuclei of each brain (A and B), likely corresponding to active enhancers (**Fig. 6A**). However, unlike the promoter ATAC-seq peak, these likely enhancer peaks did not survive the IDR analysis (**Fig. 6A**), implying that this analysis may be too stringent for the postmortem human brain tissue. These data are also consistent with the quality control genome distribution data indicating that the promoter peaks are mainly retained but the enhancer peaks are lost following the IDR analysis (**Fig. 5B)**.

For *GFAP*, consistent with this gene being a glial marker, we found chromatin to be open in non-neuronal (NeuN-) cells but closed in neuronal (NeuN+) cells (**Fig. 6B**). In particular, we detected an ATAC-seq peak that was around the TSS, corresponded with the H3K4me3 and H3K27ac peaks in the Roadmap Epigenomics datasets, and is thus consistent with an active promoter in non-neuronal (NeuN-) cells (**Fig. 6B**). The other two ATAC-seq peaks, both within 5 kb upstream of the TSS, are likely to represent enhancers, as also implied by the combination of H3K27ac and H3K4me1 in these regions (**Fig. 6B**). Strikingly, using the whole genome bisulfite sequencing (WGBS) method, we show that all three *GFAP* regulatory regions (promoter and two enhancers) which show differential chromatin accessibility in non-neuronal vs. neuronal cells, also show differential DNA methylation. In all three regions, CpG methylation levels are much higher in neuronal than in non-neuronal cells, and we validate these data by performing the bisulfite pyrosequencing analysis of the three CpG sites located in the proximal enhancer (**Fig. 6B**). Another finding worth noting is that the Roadmap Epigenomics data show both active histone marks (H3K4me3 and H3K27ac) and repressive histone mark (H3K27me3) within the *GFAP* regulatory regions (**Fig. 6B**), obviously due to the fact that the data are generated from bulk brain tissue, containing both neuronal and non-neuronal cells. With our cell type specific epigenomics assays, we show that we can clearly distinguish between neuronal and non-neuronal cells using both ATAC-seq and WGBS assays. These data further show that, although the postmortem human brain is a more challenging tissue to perform the cell-type specific chromatin accessibility assay, the ATAC-seq method is generating reliable data.

## DISCUSSION

Our study shows that the cryopreservation method impacts the number of open chromatin regions that can be identified from frozen brain tissue using the cell-type specific ATAC-seq assay. However, we also show that all three common freezing procedures that we tested generate consistent and robust ATAC-seq data and enable the identification of functional regulatory elements, promoters and enhancers, in neuronal cells. Our study also implies that the broad biological interpretation of chromatin accessibility data is not significantly affected by the freezing condition.

It is worth noting that all well-established epigenomics assays, including ATAC-seq, were initially developed and still most commonly used with cell culture systems^16^, in situations when there is easy access to millions of freshly harvested cells. Many quality standards, such as those employed by the ENCODE project, were also developed based on the data produced from cell lines^10^. Working with tissue is more complicated, particularly in neuroscience, where working with frozen brain tissue is almost unavoidable. Many study designs, whether those including birth cohorts when studying early life stressors or adult exposures to drug or stress, will involve large numbers of animals that need to be processed for molecular analyses. Working with fresh brains under these circumstances is usually time-prohibitive. However, there is a possibility to use a different cryopreservation technique, which can have a big impact on downstream genomic results, but this is rarely optimized.

We decided to test different freezing conditions for the ATAC-seq particularly, because this method was initially thought to be incompatible with frozen tissue^16^. In fact, there were studies which showed that regular flash freezing of neuronal cells barely produce any ATAC-seq peaks, whereas the enrichment is highly improved by using a slow freezing protocol with a cryoprotective medium containing dimethyl sulfoxide (DMSO)^18^. Therefore, we used the slow freezing protocol (MED condition) as a reference in our study, although we were aware that even this protocol would be difficult to implement in our study designs when tens of animals need to be sacrificed in a short period of time. The other two protocols that we included are typically used in our laboratory. Flash freezing with liquid nitrogen (LN2 condition) is typically utilized for studies that are medium-size and for brain regions that are easier to dissect from fresh brains (e.g. ventral/dorsal hippocampus, nucleus accumbens). On the other hand, for larger studies, it is very beneficial to quickly submerge a whole freshly extracted brain into hexane cooled on dry ice (HEX condition); this allows for fast cooling while preserving the structure of the brain that would be easier to dissect on a later occasion.

While the MED condition was, indeed, the most efficient method and produced the highest number of peaks and the peaks that were strongest, it is striking that the data looked very consistent among the three conditions. It is also interesting that the HEX condition was more efficient than the LN2 condition, suggesting a positive effect of hexane on preserving chromatin structure. Based on our data, looking at specific genes and more generally, all three methods are producing high quality data and can be successfully used for neuroepigenomic analyses. It is obvious, though, that having more peaks is going to provide useful additional information, particularly because these small peaks found in the MED condition are likely to correspond to cell-type specific enhancers^13^. However, we should keep in mind that we had only about 40 million paired-end reads per sample in all conditions; hence, there is a possibility that simply increasing the number of reads may uncover those MED-specific peaks in other two conditions. In addition, if these “weak” peaks correspond to a very small population of neuronal cells, developing a more neuronal subtype specific analysis would be likely beneficial to learn more about these functional elements.

Finally, we also explored the cell-type specific ATAC-seq analysis in *post mortem* human brain. Although comparable studies in human brain have been published^22-26^, no one has compared the analyses of mouse and human brain side by side. While we observe that the quality of data decreases in the human brain, by looking at FRiP scores in other human brain studies, we can also see that these quality metrics can vary widely^22^, which is always a challenge when working with *post mortem* tissue involving many variables (PIM, pH value, etc.). We also note that many non-promoter peaks are lost after the IDR analysis of human ATAC-seq data, implying that the same analytical methods may not be appropriate for the mouse and human brain chromatin accessibility analysis. It is, therefore, not surprising that the published studies pooled the data from different subjects to call the ATAC-seq peaks or merged individual peaks across all samples^22,26^. While the pooling approach prohibits accounting for individual variability, it gives more power to detect weaker peaks that may be lost by too stringent analysis that is appropriate for mouse brain tissue. Overall, our data confirm the challenges of doing chromatin analysis on *post mortem* human brain tissue but also shows that these studies are revealing important information about gene regulation in the human brain. We show that chromatin analysis is able to clearly mark cell-type specific gene regulatory elements, consistent with the (presumably more stable) DNA methylation signature, in the human brain.

In summary, the ATAC-seq coupled with FANS is a powerful method to capture cell-type specific chromatin accessibility information in the mouse and human brain. Our study provides alternative brain preservation methods that generate high quality ATAC-seq data while fitting in different study designs, and further encourages the use of this method to uncover the role of epigenetic (dys)regulation in healthy and malfunctioning brain.

## METHODS

### Mouse brain tissue samples

Nine-week-old C57BL/6J male mice (n=6) were purchased from Jackson Laboratory. Mice were housed in groups of three per cage, were given *ad libitum* access to food and water, and were kept on a 12:12h light:dark cycle with lights on at 8 am. After two weeks of habituation, mice were sacrificed an hour after the beginning of the light phase. Each brain was immediately removed from the skull and three methods were used to preserve the cortical tissue (n=2 per freezing condition, **Figure 1A**): 1) *Slow freezing in freezing medium (MED)* — frontal cortices were bilaterally dissected on ice and immediately submerged in a cryopreservation medium containing 90% of fetal bovine serum (Gibco™ A3160601) and 10% DMSO (Sigma-Aldrich, D8418-50ML). Samples were then frozen in an isopropyl alcohol-containing Mr. Frosty chamber (ThermoFisher Scientific) at −80°C, at a rate of cooling of −1°C/minute, as previously described^18^; 2) *Flash-freezing in dry ice-cooled hexane (HEX)*—whole brain was submerged in dry ice cooled hexane (−95°C) for 2 minutes and then stored at −80°C. Frontal cortices were bilaterally dissected the next day at −20°C and stored at −80°C until nuclei prep; 3) *Flash-freezing in liquid nitrogen (LN2)* — frontal cortices were bilaterally dissected on ice, immediately frozen in liquid nitrogen (−195°C), and then stored at −80°C. All animal procedures were in full compliance with the National Guidelines on the Care and Use of Laboratory Animals and were approved by the Institutional Animal Care and Use Committee at Fordham University.

### Human post mortem brain tissue samples

100 mg of the pulverized anterior cingulate cortex (ACC) tissue from 2 individuals, 49-year-old African American male (Brain A, PMI = 15.5) and a 52-year-old Caucasian male (Brain B, PMI=23), was obtained from the NIMH Human Brain Collection Core. At the time of death, the subjects had no history of psychiatric disorder, were non-smokers, and tested negatively for nicotine, cotinine, ethanol, THC, cocaine, and opiates. There was also no evidence of the use of psychiatric medications, including benzodiazepines, mood stabilizers, or antidepressants. Specimen information is presented in **Supplementary Table S6**.

### Fluorescence-activated Nuclei Sorting (FANS)

The isolation and purification of neuronal nuclei from bulk brain tissue was performed with a protocol which we previously published^15^. Briefly, dissected bilateral cortical mouse tissue or 100 mg of *post mortem* human ACC tissue was homogenized using a tissue douncer. Nuclei were then extracted through a sucrose gradient using ultracentrifugation at 24,400rpm (101,814.1 rcf) for 1 hour at 4°C in a Beckman ultracentrifuge. Following ultracentrifugation, nuclei were resuspended in Dulbecco’s phosphate-buffered saline (DPBS) and incubated with a mouse monoclonal antibody against NeuN, a neuronal nuclear protein, conjugated to AlexaFluor-488 (1:1000; Millipore, MAB 377X). Just prior to sorting, nuclei were also incubated with DAPI (1:1000; ThermoFisher Scientific, 62248) and then filtered through a 35-µm cell strainer.

Sorting was performed at the Albert Einstein College of Medicine Flow Cytometry Core Facility using the FACSAria instrument (BD Sciences). To ensure proper gating, three controls were used: i) IgG1 isotype control-AlexaFluor-488, ii) DAPI only, and iii) NeuN-AlexaFluor-488 only. Using data acquired from these controls, gating was adjusted for each sample so that single nuclei were selected from debris and any clumped nuclei, and so that the NeuN+ (neuronal) population could be sorted and separated from the NeuN- (non-neuronal) population (**Supplementary Fig. 1**). From each mouse sample we collected 75,000 NeuN+ nuclei in tubes pre-coated with BSA containing 200µl of DPBS. From each human sample we collected 75,000 NeuN+ and 75,000 NeuN-nuclei for ATAC-seq. An additional 83,000 NeuN+ and 149,000 NeuN-nuclei were collected from Brain A for whole-genome bisulfite sequencing.

### ATAC-Seq Assay

Following sorting, 75,000 cortical, neuronal (mouse, human) and non-neuronal (human) nuclei were centrifuged at 5,557rpm (2,900rcf) for 10 minutes at 4°C. After removal of the supernatant, the nuclei-containing pellet was resuspended with 50 µl of the transposase reaction mix which contained 25 µl of the 2xTD reaction buffer and 2.5 µl of Tn5 Transposase (Nextera DNA Library Preparation Kit, Illumina, FC-121-1030; note: these reagents are now available in the Illumina Tagment DNA TDE1 Enzyme and Buffer Kit). Transposition was performed for 30 minutes at 37°C after which the transposed DNA was purified using MinElute PCR Purification Kit (Qiagen, 28004) and eluted in 10 µl of elution buffer (EB) and stored at −20°C.

Continuing with the preparation of sequencing libraries, indexing and amplification of the transposed DNA samples was performed by combining 10 µL of transposed DNA with the following: 5 µL of the Nextera i5 and i7 indexed amplification primers (Nextera Index Kit, Illumina, FC-121-1011), 25 μl of the NEBNext High-Fidelity 2x PCR Master Mix (New England Biolabs, M0541S), and 5 µL of the Illumina PCR Primer Cocktail (PPC, Illumina FC-121-1030). PCR conditions were as follows: an initial cycle of 5 minutes at 72°C then 30 seconds at 98°C; followed by 5 cycles of 10 seconds at 98°C, 30 seconds at 63°C, and 1 minute at 72°C. Following PCR, samples were held at 4°C. To determine the remaining number of PCR cycles needed for each sample, we performed a 20-cycle qPCR reaction with the following components: 5 µL of PCR-amplified DNA, 5 μl NEBNext High-Fidelity 2x PCR Master Mix, 0.5 µL Illumina PCR Primer Cocktail and 4.5 µL of 0.6x 2xSYBR green I (Invitrogen, S7563). From the results of this qPCR we determined that each library required 4-5 additional PCR cycles in order to achieve optimal amplification without compromising library complexity. Following amplification, purification of the libraries was performed using the *MinElute PCR Purification Kit* and elution was performed with 20 µL of the elution buffer. To assess library quality, we used 2.2 % agarose gel electrophoresis (FlashGel DNA System, Lonza) and Bioanalyzer High-Sensitivity DNA Assay (**Supplementary Figure 2**). Prior to sequencing, quantification of ATAC-seq libraries was performed using the Qubit HS DNA kit (Life Technologies, Q32851) and the qPCR method (KAPA Biosystems, KK4873). Sequencing of 100-bp paired-end reads was performed on Illumina HiSeq 2500 instruments at the Albert Einstein College of Medicine Epigenomics Shared Facility.

### ATAC-seq Data Analysis

First, adapters were trimmed and reads were aligned to the appropriate reference genome (mm10 for mouse, hg38 for human) using BWA-MEM software^47^. Peak calling was performed using MACS2 software on reads uniquely aligned to the corresponding genome as previously described^16^. The number of aligned reads in mouse samples ranged from 42.3 to 52.7 million. Thus, we performed analyses with and without down-sampling (**Supplementary Table 1**). The down-sampling factors were calculated to have a comparable number of reads-in-peaks in all samples. For each sample, the number of aligned reads and percentage of reads in peaks (%RiP) before and after downsampling are shown in **Supplementary Table 1**. Read numbers, %RiP values, feature distribution plots, and the principal component plot were obtained using the ChIPQC Bioconductor package^48^. We used results after the down-sampling for following analyses. For each group having two biological replicates, we selected high confidence peaks for further analysis by calculating the irreproducible discovery rate (IDR) between replicates and continuing with the peaks passing a cutoff of IDR < 0.05^49^. The number of peaks for each sample and number of IDR-peaks for each group is shown in **Supplementary Table 1**. Annotation of peaks and the overlap status of peaks between groups was analyzed using the *annotatePeakInBatch* and *findOverlapsOfPeaks* functions, respectively, in the ChIPPeakAnno Bioconductor package^50^. Gene Ontology and KEGG Pathway analyses were performed on annotated peaks using the clusterProfiler Bioconductor package^51^. HOMER motif analysis was performed using the findMotifsGenome.pl function of the HOMER software (v3.12)^52^.

### Whole-genome Bisulfite Sequencing

A subset of nuclei isolated from a postmortem human ACC sample (Brain A) were used to extract genomic DNA for DNA methylation analysis. Following sorting, 83,000 neuronal and 149,000 non-neuronal human nuclei were re-pelleted by adding DPBS, sucrose buffer (1.8 M Sucrose, 3 mM Mg(Ace)2, 10 mM Tris-HCl (pH 8.0), and 1 mM DTT), 1M CaCl2, and 1M Mg(Ace)2, then leaving on ice for 15 minutes. Afterwards, the samples were centrifuged at 4,000 rpm (3094.6 rcf) for 15 min at 4°C and the supernatant was aspirated leaving the nuclei pellet. The pellet was then resuspended in DNA prep buffer (100mM NaCl,10mM Tris (pH 8.0), 10mM EDTA, and 0.5% SDS). Protein degradation was performed by adding Proteinase K (Zymo) and Qiagen Protease (Qiagen), then incubating the samples at 50°C for 2 hours. Phenol-chloroform extraction was then performed using phenol:chloroform:isoamyl alcohol (25:24:1). Following extraction, the aqueous phase was transferred to a new tube and 5M NaCl, 100% ethanol, and GlycoBlue were added before placing the samples overnight at −20°C. Samples were then centrifuged at 14,000 rpm (18406.7 rcf) for 30min at 4°C. After aspirating the supernatant, the pellet was washed with 75% ethanol and then centrifuged at 14,000 rpm (18406.7 rcf) for 5min at 4°C. The supernatant was aspirated, and any remaining ethanol was allowed to evaporate before adding elution buffer. DNA concentration was measured using the Qubit HS DNA kit and DNA was stored at −20°C. For the whole genome bisulfite sequencing (WGBS) method, DNA library preparation and sequencing were performed at the New York Genome Center based on our previously established transposase-based library preparation protocol optimized for the Illumina HiSeq X platform^53^.

### WGBS Data Analysis

For WGBS data analysis, reads were first adaptor-trimmed with cutadapt v1.9.1^54^. Alignment was performed against the GRCh38 human reference genome using bwa-meth with R1 reads mapped as a G>A converted read and R2 reads mapped as a C>T converted R1 read^55^. Read alignments were marked for duplicates using Picard v2.4.1 in a strand-specific manner based on the orientation of the read with the reference genome. Due to an artifact related to our end-repair protocol, we used a custom Perl script called filterFillIn2, which allowed us to mark and exclude reads with four consecutive methylated CHs beyond the first nine bases of the read. The bedGraph file with methylation ratios was generated by extracting methylation ratios from the alignment files using Bismark (http://www.bioinformatics.babraham.ac.uk/projects/bismark/).

### Bisulfite Pyrosequencing

Methylation levels at three CpG sites in the *GFAP* gene were validated using the bisulfite-pyrosequencing method. Briefly, 20 ng of genomic DNA derived from sorted neuronal and non-neuronal nuclei of a postmortem human ACC sample (Brain A) was used for bisulfite conversion (EpiTect Bisulfite Kit, Qiagen). For each sample, 10 ng of bisulfite converted DNA was run in a PCR using the PyroMark PCR kit (Qiagen) and the following PCR primers: forward primer (5’ – TTTTTTTTGGGTATAGGTTGAATAGAGTT -3’) and biotinylated reverse primer (5’ –Biosg-AACCCCCTAAAAACTACCCTTACTA – 3’). Biotinylated PCR products were processed on a vacuum workstation and pyrosequencing was performed on an Advanced PyroMark Q24 pyrosequencer using the advanced CpG reagents (Qiagen) and the sequencing primer 5’ – AGGTTGAATAGAGTTTTGT -3’). Before sample analysis, the pyrosequencing assay was validated using a standard curve by analyzing 0, 20, 40, 60, 80, and 100% methylated human genomic DNA standards. Those standards were generated by mixing commercially available unmethylated and hypermethylated DNA standards (EpiTect PCR Control DNA Set, Qiagen) resulting in 10 ng of total bisulfite-converted DNA. Methylation levels of single CpG sites were quantified using PyroMark Q24 Advanced software (Qiagen).

### Public Data Used

We used the following ChIP-seq datasets from the mouse ENCODE project (https://www.ncbi.nlm.nih.gov/bioproject/PRJNA63471), for postnatal zero (0) day C57BL/6 mouse forebrain: ENCSR258YWW (H3K4me3), ENCSR094TTT (H3K27ac), ENCSR465PLB (H3K4me1), ENCSR070MOK (H3K27me3), and ChromHMM PO_forebrain_15_segments (http://enhancer.sdsc.edu/enhancer_export/ENCODE/chromHMM/pooled/tracks/). For the human study, we used the following dataset from the NIH Roadmap Epigenomics Project (https://www.ncbi.nlm.nih.gov/bioproject/PRJNA34535), for *Homo sapiens* cingulate gyrus male adult (81 years): ENCSR032BMQ (H3K4me3), ENCSR355UYP (H3K27ac), ENCSR443NEP (H3K4me1), and ENCSR022LOH (H3K27me3).

## Acknowledgement

This study was supported by the NARSAD Young Investigator Grant from the Brain & Behavior Research Foundation to M.K. We would further like to acknowledge the resources of the Center for Epigenomics at the Albert Einstein College of Medicine. The postmortem human brain tissue used in this research was obtained from the Human Brain Collection Core, Intramural Research Program, NIMH (http://www.nimh.nih.gov/hbcc); we extend our thanks to Drs. Barbara Lipska and Stefano Marenco for providing the tissue. Finally, we would like to thank Dr. Shahina Maqbool for her assistance with massively parallel sequencing.

## Author Contributions

M.S. and M.K. designed the study; I.J. and M.K. performed experiments; L.T. performed nuclei sorting; D.R. and M.S. performed bioinformatics analysis; J.M.G. contributed computational resources; D.R., M.S., and M.K. interpreted the data and constructed the figures; D.R. and M.K. wrote the manuscript with input from all authors.

