## Supplementary Material for "Cell Type-Specific Chromatin Accessibility Analysis in the Mouse and Human Brain"

### Supplementary Material - Table of Contents


**Supplementary Figures:**

**Supplementary Figure 1.** Separation of neuronal nuclei using fluorescence-activated nuclei sorting (FANS)

**Supplementary Figure 2.** ATAC-seq library quality control

**Supplementary Tables**

**Supplementary Table 1: Mouse brain samples and ATAC-seq data basic information** (uploaded as a separate excel file):

**Supplementary Table 2: Mouse ATAC-seq IDR peaks for each condition** (uploaded as a separate excel file):

Suppl. Table 2a: MED total IDR peaks

Suppl. Table 2b: HEX total IDR peaks

Suppl. Table 2c: LN2 total IDR peaks

**Supplementary Table 3: Mouse significantly enriched GO terms** (uploaded as a separate excel file):

Suppl. Table 3a: MED – GO:Biological Process (BP) enriched terms

Suppl. Table 3b: HEX – GO:Biological Process (BP) enriched terms

Suppl. Table 3c: LN2 – GO:Biological Process (BP) enriched terms

Suppl. Table 3d: MED – GO:Cellular Compartment (CC) enriched terms

Suppl. Table 3e: HEX – GO:Cellular Compartment (CC) enriched terms

Suppl. Table 3f: LN2 – GO:Cellular Compartment (CC) enriched terms

Suppl. Table 3g: MED – GO:Molecular Function (MF) enriched terms

Suppl. Table 3h: HEX – GO:Molecular Function (MF) enriched terms

Suppl. Table 3i: LN2 – GO:Molecular Function (MF) enriched terms

**Supplementary Table 4: Mouse significantly enriched KEGG pathways** (uploaded as a separate excel file):

Suppl. Table 4a: MED – KEGG enriched pathways

Suppl. Table 4b: HEX – KEGG enriched pathways

Suppl. Table 4c: LN2 – KEGG enriched pathways

**Supplementary Table 5: Mouse HOMER motif analysis** (uploaded as a separate excel file):

Suppl. Table 5a: MED – HOMER motifs

Suppl. Table 5b: HEX – HOMER motifs

Suppl. Table 5c: LN2 – HOMER motifs

**Supplementary Table 6: Human postmortem brain tissue and epigenomic data basic information** (uploaded as a separate excel file)

**Supplementary Table 7: Human significantly enriched GO terms** (uploaded as a separate excel file):

Suppl. Table 7a: AP – GO:Biological Process (BP) enriched terms

Suppl. Table 7b: AP – GO:Cellular Compartment (CC) enriched terms

Suppl. Table 7c: AP – GO:Molecular Function (MF) enriched terms

**Supplementary Table 8: Human significantly enriched KEGG Pathways** (uploaded as a separate excel file):

**Supplementary Table 9: Human HOMER motif analysis** (uploaded as a separate excel file):

### Supplementary Figure 1

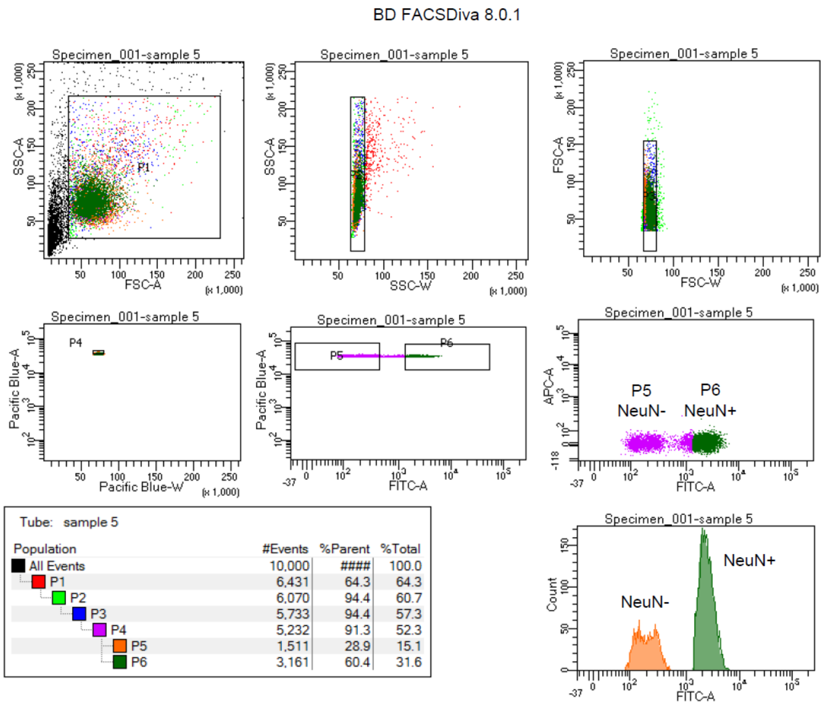


**Purification of neuronal nuclei using fluorescence-activated nuclei sorting (FANS).** Representative FANS report shows the gating strategy to: a) select nuclei from debris (P1-P3); b) ensure single nuclear sorting (using DAPI; P3); and c) select the NeuN- (non-neuronal; P5) and NeuN+ (neuronal; P6) nuclei populations. For each experiment, the gates were initially set up using two negative controls - DAPI only and isotype control + DAPI - processed without NeuN antibody; positive control containing NeuN antibody only; and each sample processed with NeuN antibody and DAPI.

**Supplementary Figure 2**

**A.**

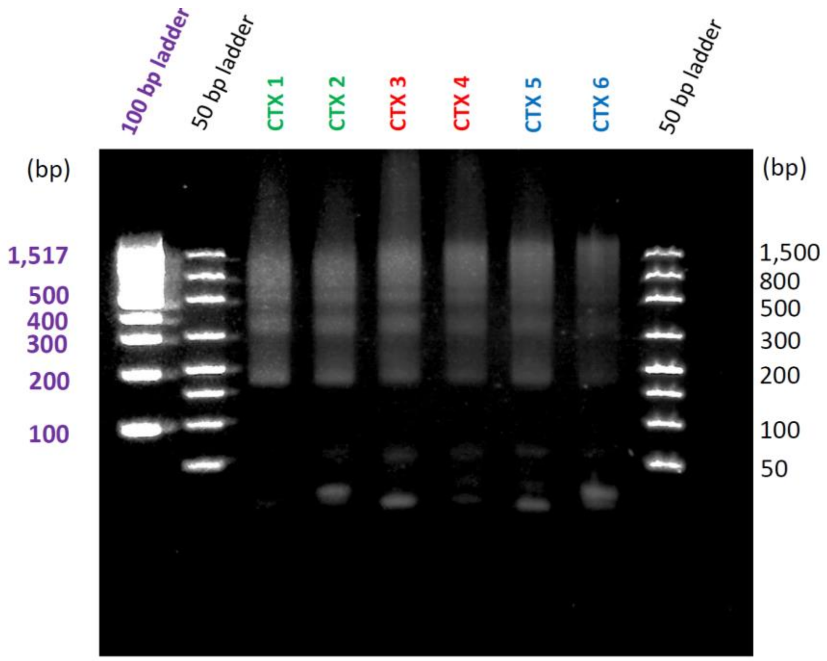


**B.**

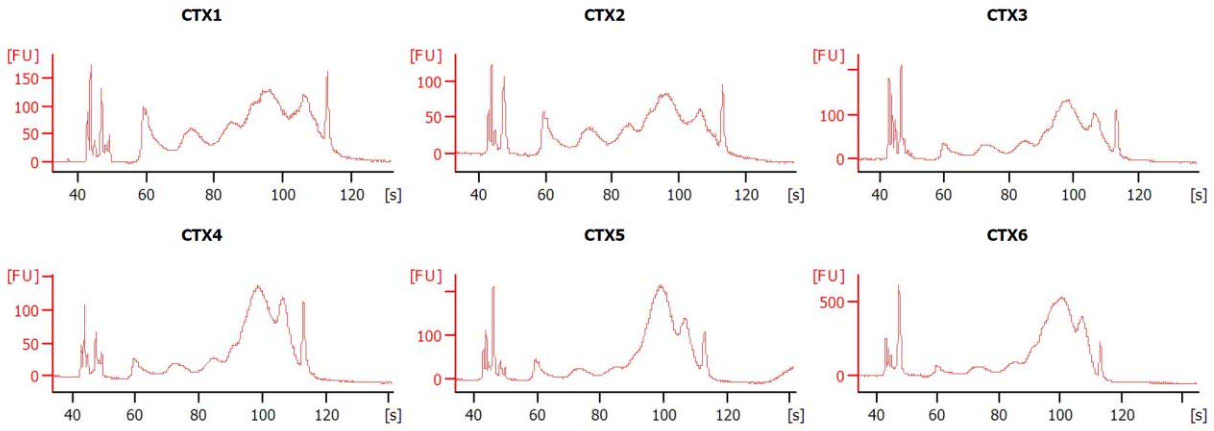


**ATAC-seq library quality control.** We assessed the quality control of our ATAC-seq libraries using: (A) 2.2 % agarose gel electrophoresis, and (B) Bioanalyzer High-Sensitivity DNA Assay. In both images, there is a clear, nucleosomal banding pattern, typical of ATAC-seq libraries. HEX (CTX1,2), LN2 (CTX3,4), and MED (CTX5,6).
